# Cerebral laterality of writing in right- and left- handers: A functional transcranial Doppler ultrasound study

**DOI:** 10.1101/2020.07.14.203588

**Authors:** Marietta Papadatou-Pastou, Panagiotis Sampanis, Ioannis Koumzis, Sofia Stefanopoulou, Dionysia Sousani, Athina Tsigkou, Nicholas A. Badcock

## Abstract

The cerebral lateralization of written language has received very limited research attention in comparison to the wealth of studies on the cerebral lateralization of oral language. The purpose of the present study was to further our understanding of written language lateralization, by elucidating the relative contribution of language and motor functions. We compared written word generation with a task that has equivalent visuomotor demands but does not include language: the repeated drawing of symbols. We assessed cerebral laterality using functional transcranial Doppler ultrasound (fTCD), a non-invasive, perfusion-sensitive neuroimaging technique in 23 left- and 31 right-handed participants. Findings suggest that the linguistic aspect of written word generation recruited more left-hemispheric areas during writing, in right-handers compared to left-handers. This difference could be explained by greater variability in cerebral laterality patterns within left-handers or the possibility that the areas subserving language in left-handers are broader than in right-handers. Another explanation is that the attentional demands of the more novel symbol copying task (compared to writing) contributed more right-hemispheric activation in right-handers, but this could not be captured in left-handers due to ceiling effects. Future work could investigate such attentional demands using both simple and complex stimuli in the copying condition.

## Introduction

An overwhelming number of studies have investigated cerebral language lateralization using overt or covert oral language production or language comprehension tasks (e.g., Papadatou-Pastou et al., 2017; Groen et al., 2012; Petit, Badcock, & Woolgar, 2020). On the contrary, very few studies have investigated the cerebral lateralization of written language (e.g., Kondyli et al., 2017) or the neural underpinnings of writing in general (e.g., Bartoň, et al., 2020; Planton et al., 2013; Rapp & Purcell, 2019; Vinci-Booher et al., 2019), although writing is starting to receive research attention (e.g., Palmis et al., 2019; Planton, Jucla, Démonet, & Soum-Favaro, 2019; Yang et al., 2019). Importantly, only a handful of studies have investigated cerebral laterality for writing comparing left- and right-handers (Kondyli, et al., 2017; Siebner et al., 2002; Zaman et al., 2002).

Writing is a skill that demands the contribution of several cognitive and visuomotor functions (for a review see Purcell et al., 2011) and is widely used in education and everyday life for communication, as well as for archiving information, ideas, and stories across time and space. Moreover, writing allows for the investigation of the brain’s capacity to acquire skills that are not biologically predetermined, given that written language is a relatively recent human invention (Purcell et al., 2011). Writing is affected in several conditions, such as Alzheimer’s disease (e.g., Hayashi et al., 2011), learning disabilities (Graham, Collins, & Rigby-Wills, 2017), schizophrenia (e.g., Tigges et al., 2000), cerebrovascular disease (e.g., Otsuki et al., 1999), and traumatic brain injury (Yorkston et al., 1997), possibly due to the complex nature of writing. Therefore, it is important to reach a clearer understanding of writing’s neural network, both when studying healthy individuals as well as when studying pathological populations. A curious facet of writing is that about 70% of left-handers (but only about 10% of right-handers) write with the hand that is ipsilateral to their (oral) language-dominant hemisphere (considering that language functions are controlled by the left hemisphere in about 90% of right-handers and 70% οf left-handers; Carey & Johnstone, 2014). This brings us to the issue of handedness.

Left-handers constitute about 10% of the population (Papadatou-Pastou et al., 2020), making it important to account for this phenotypic variation if we are to understand human brain function. Left-handers have in fact been recognized as a compelling and widely available, but largely untapped, resource for neuroscience studies (Bailey et al., 2020; Willems, Van der Haegen, Fisher, & Francks, 2014). As mentioned above, left- and right-handers differ in the cerebral organization of oral language functions (e.g., Knecht, Dräger et al., 2000), although handedness is a weak indicator of cerebral lateralization for language, accounting for only 8-16% of the variance in cerebral lateralization (Groen et al., 2013). Strongly atypical individuals in terms of hemispheric lateralization are more likely to be left-handed, although individuals with typical and bilateral dominance can also be left-handed (Mazoyer et al., 2014). Further, it has been shown that stronger left-hand preference is linked to a higher chance of atypical language lateralization (Somers, Aukes et al., 2015). Given (i) the link between handedness and cerebral lateralization for oral language, (ii) that the majority of left-handers write with the hand ipsilateral to their language-dominant hemisphere, and (iii) that the motor component of writing activates different hemispheres in left- and right-handers, it follows that cerebral lateralization for writing could differ between left- and right-handers.

The neural underpinnings of writing were initially studied in neurological patients. Lesion studies showed that impaired writing can result from localized brain damage in the superior parietal lobe, supramarginal gyrus, angular gyrus, Wernicke’s area, or Broca’s area (Roeltgen, 1993; Beeson et al., 2003). While lesion studies continue to contribute to our understanding of the neural underpinnings of writing (e.g., Tao & Rapp, 2019), in the last two decades neuroimaging (functional Magnetic Resonance Imaging [fMRI] and Positron Emission Tomography [PET]) studies have investigated the neural substrates of writing in the healthy human brain. A meta-analysis of 18 fMRI and PET studies (Planton et al., 2013) suggested that the core network of writing consists of a wide network of both cortical and subcortical cerebral regions, comprising of primarily writing-specific areas (left superior frontal sulcus/middle frontal gyrus area, left intraparietal sulcus/superior parietal area, and right cerebellum), non-specific motor areas (primary motor and sensorimotor cortex, supplementary motor area, thalamus, and putamen), and areas related to linguistic processes (ventral premotor cortex and posterior/inferior temporal cortex).

More recent study, Planton et al. (2017) found that the same ‘writing specific’ networks were recruited for both handwriting and drawing, with the only distinctive feature of handwriting, as opposed to drawing, being the left lateralization of the graphemic/motor frontal area (GMFA), a subpart of the superior premotor cortex. The striatum was shown to have a role in integrating stored letter-shape information with motor planning and execution during handwriting (Bartoň et al., 2020). Furthermore, the left parietal lobule relates to the sequential execution of writing, along with the left premotor cortex, the sensorimotor cortex, and the supplementary motor area (Karimpoor et al., 2018; Menon & Desmond, 2001). Writing (versus drawing) was further found to increase left-sided activation in the dorsal and ventral premotor cortex, Broca’s area, pre-supplementary motor area and posterior middle and inferior temporal gyri, without parietal activation (Potgieser et al., 2015). Bilateral activity in the cerebellum has also been observed (Katanoda et al., 2011; Karimpoor et al., 2018; Segal & Petrides, 2012; Yang et al., 2019) and is considered indicative of the representation of finger movements (left cerebellar activation) and of the coordinated movement of the index finger, in contrast to simple movements (right cerebellar activation).

However, the studies described above have investigated the neural underpinnings of writing only in right-handers. To date, merely three studies have included left-handers (Kondyli et al., 2017; Siebner et al., 2002; Zaman et al., 2002; Of note, in Golestanirad et al., 2015, three out of the 12 participants were left-handed, but they were left-lateralized for language and their results were merged with the rest of the participants). In a one-page brief article, Zaman et al. (2002) report an investigation of normal and mirror writing with dominant and non-dominant hands (four conditions: writing with the right hand, writing with the left hand, mirror-writing with the right hand, and mirror-writing with the left hand). They showed that when writing with the dominant hand, the left sensory-motor cortex and right cerebellum were activated for right-handers for normal writing, but the right sensory-motor cortex and the left cerebellum for left-handers. In the case of mirror writing, the activation was bilateral in both groups regardless of whether they were using the dominant or non-dominant hand. Siebner et al. (2002) showed that, during writing, right-handers had left-hemispheric lateralization with activation of parietal and premotor association areas; converted left-handers had a more bilateral activation pattern, including the premotor, parietal and temporal cortex; while left-handers had strong right-hemispheric lateralization. Siebner et al. (2002) also showed a graded increase in the activation of the right anterior supramarginal gyrus with the degree of left-handedness.

Neither Zaman et al. (2002) nor Siebner et al. (2002) compared writing with oral language or a non-linguistic motor activity. Zaman et al. (2002) examined writing with dominant versus non-dominant hands and Siebner et al. (2002) examined the effects of switching writing hands at a young age. More recently, Kondyli et al. (2017) compared (silent) oral language production to written language production in left- and right-handers. They found that during written word production, the degree of left-hemispheric lateralization was significantly increased for right-handers compared to silent word production, while on average left-handers presented left-hemispheric lateralization during silent word production, but right-hemispheric lateralization during writing. They suggested that a broader network of right-hemispheric areas could be used during writing in left-handers. This broader network could support motor and/or linguistic aspects of writing. However, the two phonemic fluency tasks employed in the Kondyli et al. (2017) study were not directly comparable, as only the writing task demanded visuomotor coordination and action, while the silent word production control task merely demanded word generation without including a visuomotor component. The silent word production task therefore acted as a language-control task, when a motor-control task would be better suited for this kind of investigation into the neural substrates of language.

When it comes to the technique used for the measurement of the neural underpinnings of written language, Zaman et al. (2002) and Siebner et al. (2002) employed PET and fMRI, respectively. These are high-cost techniques, prohibitive in terms of gathering large sample sizes. Therefore, PET and fMRI studies are commonly underpowered to reveal the effects of interest. As an illustration, Zaman et al. (2002) included only 12 left-handed participants (as well as 12 right-handed participants) and Siebner et al. (2002) only 6 left-handed participants (and also 11 right-handers and 11 ‘converted’ left-handers: the latter group comprising of adults who were innately left-handed for writing but were forced to use their right hand as children and became proficient right-hand writers). In Kondyli et al. (2017), the sample was considerably larger (30 left-handers and 30 right-handers) as cerebral lateralization was assessed using functional transcranial Doppler ultrasonography (fTCD). FTCD was also used in the present study.

FTCD is an efficient and reliable alternative to fMRI for the study of functional cerebral lateralization (Bishop, Watt, & Papadatou-Pastou, 2009). It is non-invasive and relatively inexpensive and is easily applied to individuals of all ages, starting from 1-year-olds (Kohler et al., 2015) up to 75-year-olds (Keage et al., 2015), in large cohorts (Knecht, Deppe et al., 2000), in longitudinal studies (Cuadrado et al., 1999), and in follow-up assessment (Lohmann et al., 2005). Results obtained with the use of fTCD are highly reproducible (Knecht, Deppe, Ringelstein, et al., 1998) and language laterality assessments performed using fTCD have very good agreement with the gold standard method in the field, the intra-carotid amobarbital procedure (Wada test; Knake et al., 2003; Knecht, Deppe, Ebner et al., 1998). FTCD and fMRI assessments of language lateralization also correlate very well (*r* = .95; Deppe et al., 2000, ρ = .75; Sommers et al., 2011). FTCD lends itself to the study of handwriting, as its signal is not disrupted by movement artifacts and no special equipment (e.g., fMRI-compatible tablets, see Golestanirad et al., 2015) is needed (Kondyli et al., 2017). In fMRI and PET studies, lateralization is usually determined by calculating the difference between the activated brain regions in the left and the right hemisphere relative to the sum of all activated regions in both hemispheres. FTCD provides comparable information in a much more efficient way, by directly comparing the relative blood flow velocity changes in the two middle cerebral arteries (MCAs). The quantitative measures obtained by fTCD are moreover not biased by variable group-defined statistical thresholds, as is often the case in the analysis of fMRI data (for a discussion see Johnstone et al., 2020). The typical sensitivity in these studies for detecting perfusion asymmetries between two basal arteries is of the order of 1% (Deppe et al., 1997a; Knecht et al., 1997, 1998a). FTCD’s spatial resolution is low, restricted to the basal artery territories, but it has excellent temporal resolution. FTCD is further limited by the fact that some individuals lack an acoustic temporal bone window for insonation of the MCAs and that blood flow changes indicate -but do not directly measure-functional cortical activation.

Returning to the issue of handedness, writing hand is the most commonly used criterion to determine handedness (Papadatou-Pastou et al., 2019). However, writing hand gives a mismatch with hand preference inventories of 13.5% for left-handers (the mismatch is negligible for right-handers: 0.4%) (Papadatou-Pastou et al., 2013). Other measures of handedness comprise either hand preference inventories (asking which hand is preferred for several everyday activities), tasks that quantify hand preference in terms of an internally consistent continuum, such as card-reaching in different locations (e.g., the Quantification of Hand Preference Test [QHPT], Bishop et al., 1996), or hand skill tasks (measuring the relative skill of the two hands). All measures assess direction (left-right) and/or degree of handedness. Direction of handedness is the measure typically reported, but it has been shown that the neural architecture supporting language functions, such as sentence comprehension, is modulated by degree of handedness (Newman et al., 2014). When it comes to hand skill, Brandler et al. (2013) identified the first gene to be statistically associated with handedness using the pegboard task, which is a hand skill task. This is opposed to the hand-preference measures that gave null results in previous large genetic screenings (Eriksson et al., 2010). Meta-analyses of handedness data suggest that research on handedness should include data on both hand preference and hand skill measures (Papadatou-Pastou et al., 2015; Papadatou-Pastou, 2016; Papadatou-Pastou et al., 2020). Using a comprehensive set of handedness assessment methods is indeed useful not only to explore their different properties, but mainly for comparison purposes with previous studies that may have used a single handedness measure.

The primary aim of the present study was to extend our understanding of the cerebral laterality of written versus oral language in left-compared to right-handers by disentangling the relative contribution of linguistic and motor functions in the cerebral laterality for writing. In that respect, written word generation was compared with a non-linguistic visuomotor task: written symbol copying – specifically the repeated writing of symbols (e.g., *, $, &). Symbol copying has equivalent visuomotor demands (precise coordination of fingers, wrist and arm movements, planning of sequential action, attention to visual landmarks, eye-hand coordination, and hand placement in space), but excludes generative language processes. Moreover, symbol copying more closely resembles the hand movements during writing in comparison to drawing. Thus, we will derive three laterality indices (LIs): one for the written word generation condition (LI_words_), one for the symbol copying condition (LI_symbols_), and one for the difference between them (LI_difference_ = LI_words_ - LI_symbols_). Thus, symbol copying will act as an active baseline to written word generation, allowing for the linguistic-specific activation to be estimated. We only tested writing with the dominant hand of each individual, to avoid confounding variables associated with differential practice between the two hands, such as effort made to write, possible discomfort in holding the pen, and writing larger and lower-quality letters with the non-dominant hand. Importantly, we were interested in the properties of natural writing (i.e., dominant handwriting) in left- and right-handers. Cerebral lateralization was assessed using fTCD and handedness was assessed using the Edinburgh Handedness Inventory (EHI; Oldfield, 1971), writing hand, Annett’s Pegboard task (Annett et al., 1979), and the QHPT (Bishop et al., 1996). We hypothesized that:

1. Left-handers will exhibit weaker typical (left) linguistic-specific lateralization for written language generation compared to right-handers. In other words, right-handers should present with a stronger left-hemispheric activation pattern when isolating the linguistic aspect of writing, similarly to oral language tasks. This hypothesis will be tested in two ways:

i. By examining whether the LI_difference_ differs from zero separately for left- and right-handers. We expect that the LI_difference_ will differ from zero for right-handers, but not for left-handers.
ii. examining whether left- and right-handers differ on the difference between written word generation and symbol copying. We expect to find a smaller LI_difference_ in left-handers compared to right-handers.

If we do find weaker lateralization in left-handers, motivated by Johnstone et al. (2021), we will explore four potential explanations. It may be the case that (1) left-handers have greater within-group variability compared to right-handers, (2) individual left-handers show greater inter-trial variability compared to right-handers, (3) there are fewer left-lateralized individuals who are left-handed compared to left-lateralized right-handers, and (4) lateralization is weaker in left-lateralized left-handers compared to left-lateralized right-handers.

A secondary aim of this study was to use a comprehensive set of handedness assessment criteria to examine the relationship to cerebral lateralization estimates. This deepens our understanding of handedness per se and further provides data that could be more readily comparable with studies that used a single handedness criterion. Based on previous work (Kondyli et al., 2017), we anticipate that hand preference measures will give higher correlations with the LI_difference_ compared to the hand skill measure.

## Materials and methods

### Participants

Fifty-four undergraduate and graduate students from the National and Kapodistrian University of Athens as well as members of the general public participated (mean age: 26.76 years, SD = 5.14, range: 19-40). Fourteen of the 20 male participants and 17 of the 34 female participants were right-handed, according to self-reported writing hand and the rest were left-handed (6 male and 17 female). Six of these participants had also taken part in the Kondyli et al. 2017 study (2 left-handed females, two left-handed males, and two right-handed males). All participants were monolingual, native speakers of the Greek language and they had normal or corrected-to-normal vision. They had never been diagnosed with dyslexia or dysgraphia, were free of neurological problems or other problems affecting the mobility and normal function of their hands, had not taken medication that could affect the central nervous system for at least six months, and did not report current use of illicit drugs or other substance abuse. Thirteen potential participants (2 male right-handers, 3 male left-handers, 3 female right-handers, and 5 female left-handers) were not included in the sample described above, because sonography failed due to inadequate ultrasonographic penetration of the skull by the ultrasound beam (19%, a rate similar to previous studies, e.g., Knake et al., 2003, who had an exclusion rate of 15% and Kondyli et al., 2017, who had an exclusion rate of 18.9%).

### Assessment of handedness

#### Edinburgh Handedness Inventory (EHI)

Hand preference was self-reported using the Greek translation of the Edinburgh Handedness Inventory (Oldfield, 1971). Participants indicated hand preference for ten activities, namely writing, drawing, throwing a ball, using scissors, using a toothbrush, holding a knife to carve meat, holding a spoon, holding a broom (upper hand), striking a match, and opening the lid of a box. Two additional activities referring to foot and eye preference were included (kicking a ball, and eye preferred for looking in case only one eye is used). The participants were instructed to imagine or recall which hand, foot, or eye they use when they perform each activity before answering a question. Possible responses included: “always left”, “usually left”, “no preference”, “usually right”, and “always right”.

A value of 0 was given to “always left” responses, 1 to “usually left” responses, 2 to “both equally” responses, 3 to “usually right” responses, and a value of 4 to “always right” responses. The total score of each participant was again divided by the maximum score (40), and multiplied by 100, with the LI ranging from 0 % (extreme left-handedness) to 100 % (extreme right-handedness). Individuals were classified as left-handers if their scores were below 50% and as right-handers if their scores were above 50%.

#### Pegboard

Annett’s pegboard task (Annett et al., 1979) was employed to measure relative hand skill. A 32 × 18 cm wooden equipment was used, which consisted of two attached wooden pieces with 10 holes drilled along their length. The distance between the two wooden pieces was 15 cm and the diameter of each hole was approximately 1.2 cm. Each peg was 7.0 cm in length and 1.0 cm in width. The task the participants were asked to carry out once seated in front of the pegboard, was to move all 10 pegs as quickly as possible from the full row to the empty row beginning on the side of the pegboard ipsilateral to the hand being used to perform the task. Trials for all participants started with the right hand and then the left and right hands alternated. The task was repeated three times for each hand. A stopwatch was used to time the participants. If a participant dropped a peg, the trial was repeated. Participants were instructed not to talk while carrying out the task, as talking might delay them.

The time that the first peg was touched by the participant until the time the last one was released was recorded for each trial (three trials for each hand). A Laterality Index (LI) was calculated using the formula: LI = [(RH-LH) / (RH+LH)]*100, where RH = mean time needed to move the pegs using the right hand and LH = mean time needed to move the pegs using the left hand. A negative score represented right-hand superiority, while a positive score represented left-hand superiority.

#### Quantification of Hand Preference Test (QHPT)

Hand preference was observed using the QHPT (Bishop et al.,1996). Seven positions were marked on a table, each at a distance of 40 cm from the midpoint of a baseline, at successive 30° intervals. Three cards were placed at each position, totaling 21 cards. The participants were asked to stand in front of the table with their arms resting at their sides and to pick up a named card and place it in a box in front of them. The order of the cards was random, but it was kept the same for all the participants. The hand chosen to pick up each card was recorded.

A value of 0 in the case that the left hand was used to place the card into the box, 1 point in case of changing hands, and 2 points when the right hand was used. The total points assigned to each participant were then divided by the maximum score (40) and multiplied by 100, in order to calculate an LI. This LI varied from 0% (extreme left-handedness) to 100% (extreme right-handedness). Individuals were classified as left-handers if their scores were below 50% and as right-handers if their scores were above 50%.

### Assessment of linguistic lateralization

#### Apparatus

A commercially available Doppler ultrasonography device (DWL Multidop T2: manufacturer, DWL Elektronische Systeme, Singen, Germany) was used to measure bilateral blood flow. Two 2-MHz transducer probes were mounted on a flexible headset and placed at the left and right temporal windows of the participants who were seated in front of a computer screen.

#### Lateralization tasks

The tasks were a modification of the phonemic fluency tasks by Kondyli et al. (2017), which were, in turn, based on word generation developed by Knecht, Deppe, Ebner et al. (1998). Each trial included: 35 seconds of rest, a cueing tone (accompanied by an event marker sent to the fTCD device) followed by a 5-second pause, a letter of the Greek alphabet or a symbol appearing on the center of the computer screen for 2.5 seconds, and a 12.5-second activity period. The cueing tone was used to help focus the attention on the upcoming task. There were 40 trials in total, as the standard word generation task (Knecht, Deppe, Ebner et al., 1998) comprises 20 trials and here two conditions were tested. The 40 trials were divided into 20 letter trials corresponding to written word generation and 20 symbol trials corresponding to symbol copying. Ten consecutive trials of each condition were alternated with ten consecutive trials of the other condition (e.g., 10 letter trials followed by 10 symbol trials, 10 letter trials, and 10 symbol trials). The order of presentation of the trials was counterbalanced across participants.

For the written word generation condition, the participants were asked to write down as many words as possible starting with the letter appearing on the screen. For the symbol copying condition, the participants had to copy as many times as possible the symbol appearing on the screen. The 20 letters from the Greek alphabet were chosen out of the total 24 included in the alphabet after a pilot procedure described in Kondyli et al. (2017), which ensured that the letters chosen allowed participants to produce the most words possible. A pilot study with 6 participants (4 male, mean age: 26.6 years, SD = 4.13, range: 18-35) was run in order to ensure feasibility of the protocol. The results are not included in the present report.

### FTCD data collection and analysis

Two transducer probes (2 MHz) attached to a flexible headband were placed at the temporal skull windows bilaterally. The right and left MCAs were insonated at the optimal depth for each participant (45-56 mm) and the angles of insonation were adjusted to obtain the maximal signal intensity. Visual stimuli (letters or symbols) were presented on a computer controlled by PsychoPy software (Neurobehavioural Systems; Peirce 2007, 2009; Peirce et al., 2019), which sent marker pulses to the Multi-Dop system to mark the start of each epoch. The spectral envelope curves of the Doppler signal were recorded with a rate of 100 Hz and stored for off-line processing.

Data were processed using DOPOSCCI 3.1.6 (Badcock, Holt, Holden, & Bishop, 2012; Badcock et al., 2018), a MATLAB-based toolbox (https://github.com/nicalbee). Left-and right-channel blood flow velocity was downsampled to 25 Hz, normalized to a mean of 100, and variability due to heartbeat was removed as described by Deppe et al. (Deppe et al., 1997b), but using a linear correction (see Badcock et al., 2018). The data were epoched from 18 secs before to 36 secs after the cueing tone, with baseline correction between -18 to 0 secs relative to the cueing tone. Following Kondyli et al. (2017), epochs containing cerebral blood flow volume values outside the range of 70% to 130% of the mean velocity or an absolute left-minus right channel difference of 20% were rejected. The remaining data were then averaged. The laterality index (LI) was calculated as the average left-minus-right channel difference within the period of interest (POI): 7-24 s after cueing. This POI was chosen because visual inspection of the overall evoked-flow plot suggested that a POI of 7-24 s after cuing included the maximum activation. Note: the average of the POI, rather than an average around the peak, results in a more-normal distribution (Woodhead et al., 2018) that is also statistically unbiased in terms of finding a significant LI (Petit et al., 2020). We calculated three LIs: LI_words_ for the written word generation condition, LI_symbols_ for the symbol copying condition and the difference between the two LIs (LI_difference_ = LI_words_ minus LI_symbols_). Following Kondyli et al. (2017), if less than 10 epochs were accepted for either the word or the symbol conditions, the participant was excluded from the sample (*n* = 8 in this case, an exclusion rate similar to Kondyli et al., 2017).

### Procedure

Upon arrival at the lab, the study was explained to the participants, and they were encouraged to ask questions. They gave their written consent but were explicitly told that they were free to leave at any time and without having to give any reason for doing so. The participants were tested individually in a quiet room. They were asked to sit in front of a computer, and they were given the option to watch the first few minutes of a movie while the probes were being placed. The words/symbols task followed. Once the fTCD data collection was completed, the participants were asked to perform the pegboard task and the QHPT and to fill in the Greek version of the EHI. Participants were debriefed after the completion of the study. The study was conducted with the approval of the local research ethics committee.

### Analysis

Statistical analyses were performed using R (R Core Team, 2020) and the following packages: haven (Wickham & Miller, 2020), dplyr (Wickham et al., 2021), e1071 (Meyer et al., 2020), ggplot2 (Wickham, 2016), tidyr (Wickham, 2021), car (Fox & Weisberg, 2019), rstatix (Kassambara, 2020), psych (Revelle, 2021), and tibble (Müller & Wickham, 2021). Individual fTCD LI categorization was based on whether 95% confidence intervals for the LI overlapped with zero: left = lower-bound greater than zero, right = upper-bound less than zero, and bilateral (or symmetrical) = overlap with zero (Bishop et al., 2009). For categorization based on the LI_difference_, the pooled standard deviation for the word and symbol tasks were used. However, due to subtracting two, relatively noisy signals, the categorization was heavily biased to bilateral organization, which we do not believe to be true. To reduce the variance, jackknife standard deviations for the word and symbol tasks were used (Robertson, 1991), which produced categorization more in line with our expectations. No variable violated the assumption of normally distributed data (Kim, 2013). Hypothesis 1 was tested in two ways: (i) via one-sample *t*-tests examining whether the LI_difference_ differed from zero of each handedness group, and (ii) via an independent sample *t*-test examining whether the LI_difference_ differed between handedness groups. No correction for multiple comparisons was used as only three tests address the hypothesis (Althouse, 2016; Rothman, 1990).

To explore the relationships between different handedness measures as well as the relationship to cerebral lateralization, Spearman correlations were run for the difference LI (LI_difference_ = LI_words_ minus LI_symbols_). Spearman (i.e., rank-based non-parametric) correlations were used to minimize the influence of extreme data points, common in fTCD lateralization data.

The exploration of a weaker LI_difference_ in left-handers included: (1) a Levene’s test of equality of variance to examine whether the within-group LI_difference_ variability differed between handedness groups, (2) an independent-samples *t*-test to examine whether individual inter-trial variability (i.e., individual pooled standard deviation of the LI_difference_) differed between groups, (3) a χ^2^ test of association to examine whether the number of left-lateralized individuals differed between groups, and (4) an independent-samples *t*-test to test whether, for these left-lateralized individuals, LI_difference_ differed between handedness groups. We did not test for sex differences, as the literature points to the direction of no differences (Kondyli et al., 2017) and the sampling was inadequate to test for effects of sex.

## Results

The event-related blood flow for each condition by handedness is presented in **Error! Reference source not found.** and the LIs are presented in **Error! Reference source not found.**. Descriptive statistics for the three LIs by handedness (writing hand) are presented in Table 1 and descriptive statistics for the handedness assessments are presented in Table 2. The evoked flow for written word generation is in line with previous work: right-handers showed a left-lateralized pattern (LI_words_ = 3.86) and left-handers showed a right-lateralized pattern (LI_words_ = -1.39). These patterns were the same for symbol copying though weaker in the right-handers (i.e., closer to zero; LI_symbols_ = 1.64), and stronger in the left-handers (i.e., more negative; LI_symbols_ = -2.06). Word minus symbol differences showed clear left-lateralization in right-handers (LI_difference_ = 2.22) and left-lateralization in left-handers (LI_difference_ = 0.67), albeit weaker (i.e., less differentiation). Cerebral lateralization categorization based on the raw LI_difference_ SD resulted in the following groupings: Left = 13 (28.26%), Right = 1 (2.17%), and Bilateral = 32 (69.57%); and using the jackknifed LI_difference_ SD resulted in the following groupings: Left = 40 (86.96%), Right = 5 (10.87%), and Bilateral = 1 (2.17%).

**Table 1.** Descriptive statistics for the functional transcranial Doppler sonography (fTCD) conditions. Handedness groups according to the writing hand.

| Condition | Handedness | N | Mean number of epochs | Mean LI | SD | Median | Range |
| --- | --- | --- | --- | --- | --- | --- | --- |
| $LI_{\text{words}}$ | L | 19 | 18 | -1.71 | 3.13 | -2.1 | (-5.97) - 5.88 |
|  | R | 27 | 18 | 3.63 | 2.84 | 3.77 | (-4.26) - 10.74 |
| $LI_{\text{symbols}}$ | L | 19 | 17.89 | -2.51 | 3.89 | -2.62 | (-7.1) - 7.14 |
|  | R | 27 | 17.85 | 1.36 | 2.7 | 1.6 | (-5.63) - 7.2 |
| $LI_{\text{difference}}$ | L | 19 | | 0.8 | 1.75 | 1.04 | (-2.87) - 3.39 |
|  | R | 27 |  | 2.27 | 1.66 | 2.16 | (-0.67) - 6.05 |

**Table 2.** Descriptive statistics for the handedness assessments by self-reported writing hand.

| Test | Handedness | N | Mean | SD | Median | Range |
| --- | --- | --- | --- | --- | --- | --- |
| Edinburgh Handedness Inventory (EHI) 5-point | L | 19 | 35.05 | 17.79 | 30 | 20 - 84 |
|  | R | 27 | 90.52 | 8.35 | 92 | 62 - 100 |
| Annett Pegboard | L | 19 | 0.04 | 0.03 | 0.04 | (-0.02) - 0.09 |
|  | R | 27 | -0.06 | 0.04 | -0.06 | (-0.2) - 0.04 |
| Quantification of Hand Preference (QHPT) | L | 19 | 31.08 | 24.33 | 33.33 | 0 - 73.81 |
|  | R | 27 | 56.97 | 30.66 | 57.14 | 0 0 - 100 |

To test Hypothesis 1, we conducted the following analyses:

i. One-sample *t*-tests against zero were conducted by condition and handedness (writing hand). All LIs were significantly left-lateralized for right-handers; words: *t*(26) = 6.64, *p* < .001, *d* = 1.28; symbols: *t*(26) = 2.61, *p* < .05, *d* = 0.5; difference: *t*(26) = 7.12, *p* < .001, *d* = 1.37; and the condition LIs were significantly right-lateralized for left-handers: words: *t*(18) = -2.39, *p* < .05, *d* = -0.55; symbols: *t*(18) = -2.81, *p* < .05, *d* = -0.6; difference, *t*(18) = 1.99, *p* = 0.062, *d* = 0.46. The LI_difference_ tests support the hypothesis that left-handers will exhibit weaker typical (left) linguistic-specific lateralization for written language generation compared to right-handers.
ii. The second test was conducted with independent sample *t*-tests on the LI_difference_ between handedness groups; Left: M = 0.80, SD = 0.4; Right: M = 2.27, SD = 0.32. Based on writing hand, right-handers had higher LI_difference_ values, with a strong effect size, also supporting the hypothesis; *t*(34) = 2.89, *p* < .001, *d* = 0.85.

Given evidence for atypical linguistic specific lateralization, we explored why this might be the case in four ways:

1. whether the group variability differed between groups. Left- and right-handers did not differ in terms of LI_difference_ variability, *F*(1, 44) = 0.093, *p* = 0.762;
2. whether individual inter-trial LI_difference_ variability differed between groups. No evidence of a difference was found, inter-trial variability, *t*(44) = -0.21, *p* = 0.836 *d* = - 0.06. Individual LI_difference_ variability was based on the pooled standard deviation from the two conditions;
3. whether the number of left-lateralized individuals differed between groups. The number of individuals categorised as left lateralized based on the LI_difference_ (jackknifed SD) was not statistically different between left- (*n* = 15, 79%) and right-handers (*n* = 25, 93%), χ(1) = 1.83, *p* = 0.176;
4. whether, for these left-lateralized individuals, LI_difference_ differed between groups. Left- and right-handers categorised as left-lateralized based on LI_difference_ did not differ in terms of LI_difference_, *t*(38) = 2.08, *p* = 0.044, *d* = -0.06; although the left-handers were numerically lower (left-handers: M = 2.7, SD = 0.73; right-handers: M = 3.4, SD = 1.54; note, the SD did not differ between groups: Levene’s *F*(1, 38) = 5.41, *p* = 0.03). Please note that these are exploratory analyses and (4) is based on a fraction of the original sample.We conducted a further analysis to understand the visual appearance of a shift in the right-channel data between conditions in the right-handers: right activation appeared to be higher in the symbol copying condition, relative to written word generation (see Figure 1). We were interested in the 3-way interaction of a 2 x 2 x 2 mixed-design ANOVA with writing hand (right or left) as the between-subjects factor and condition (words or symbols) and channel (left or right) as within-subjects factors. The interaction was statistically significant, *F*(1, 44) = 8.41, *p* < .01, η_p_^2^ = 0.161. Splitting the data by handedness, the interaction for follow up 2 (condition) x 2 (channel) repeated-measures ANOVAs was significant for right-handers, *F*(1, 26) = 50.82, *p* < .001, η_p_^2^ = 0.662, but not for left-handers, *F*(1, 18) = 3.96, *p* = 0.062, η_p_^2^ = 0.180. This supports observations that right activation was higher for symbol copying (versus written word generation) for right-handers.

**Figure 1:**
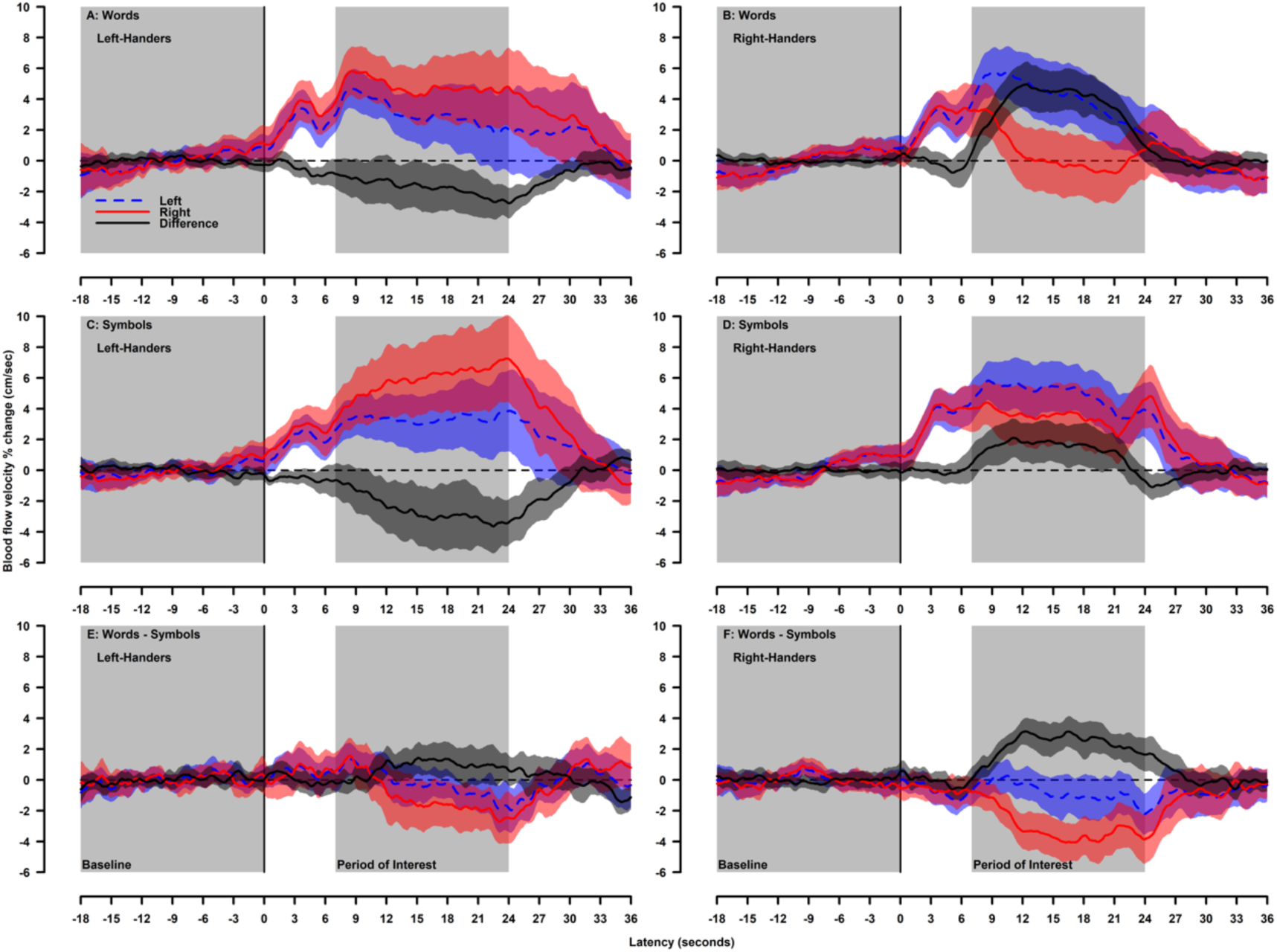
Event-related blood flow for condition (written word generation = top row, symbol copying = second row, written word generation minus symbol copying = bottom row) by handedness (left-handers = left column, right-handers = right column). Left (blue dash), right (red), and left minus right difference (black) channels are depicted with their respective 95% confidence interval. Analysis baseline and period of interest timing are depicted by grey columns. Handedness groups according to the writing hand.

**Figure 2:**
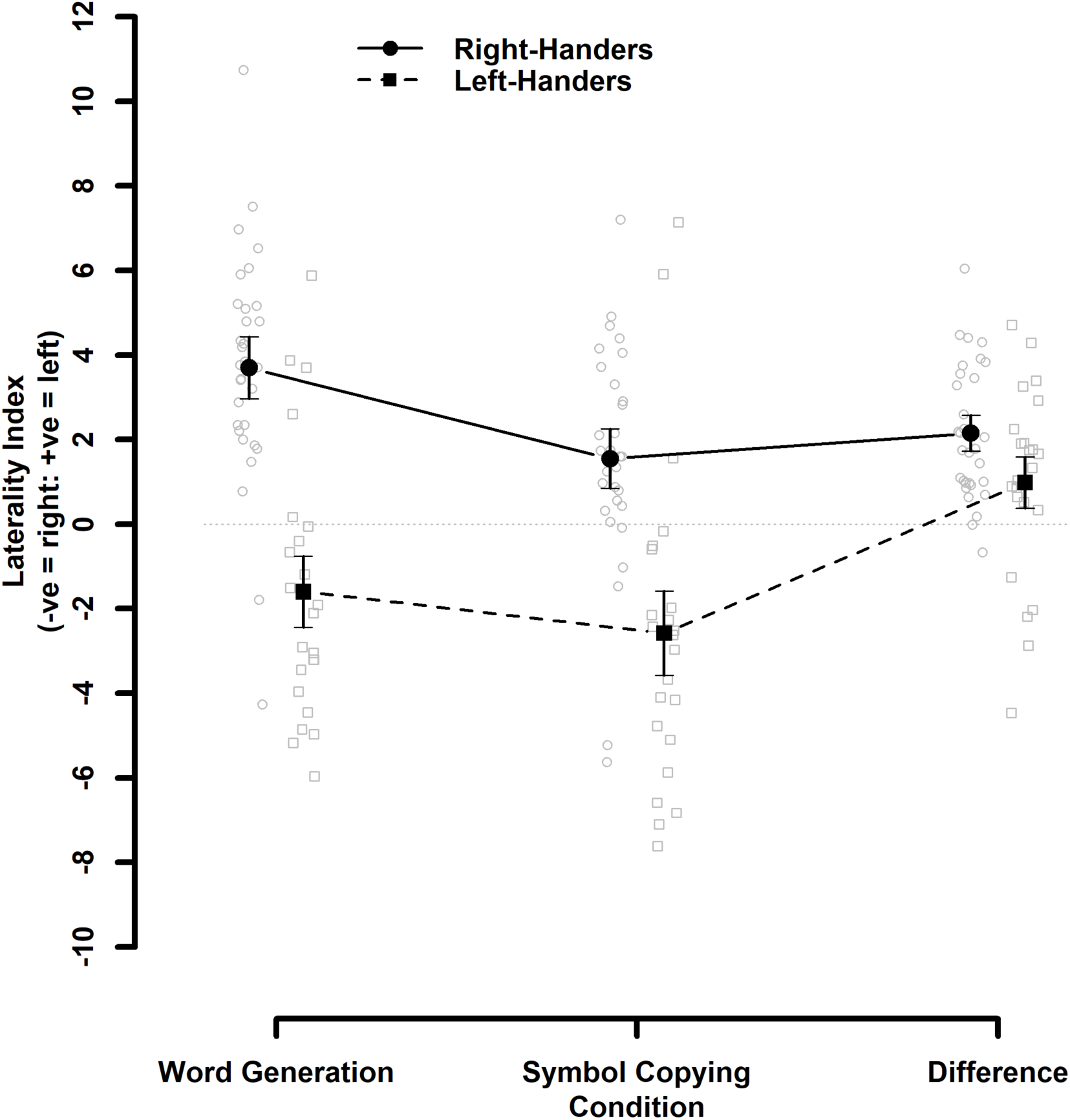
Group laterality indices (LIs) for condition (written word generation, symbol copying, written word generation minus symbol copying) by handedness (right-handers and left-handers denoted by circles with solid line and squares with broken line, respectively). Error bars represent the 95% confidence intervals. Handedness groups according to the writing hand. Grey symbols represent individual data points.

**Figure 3:**
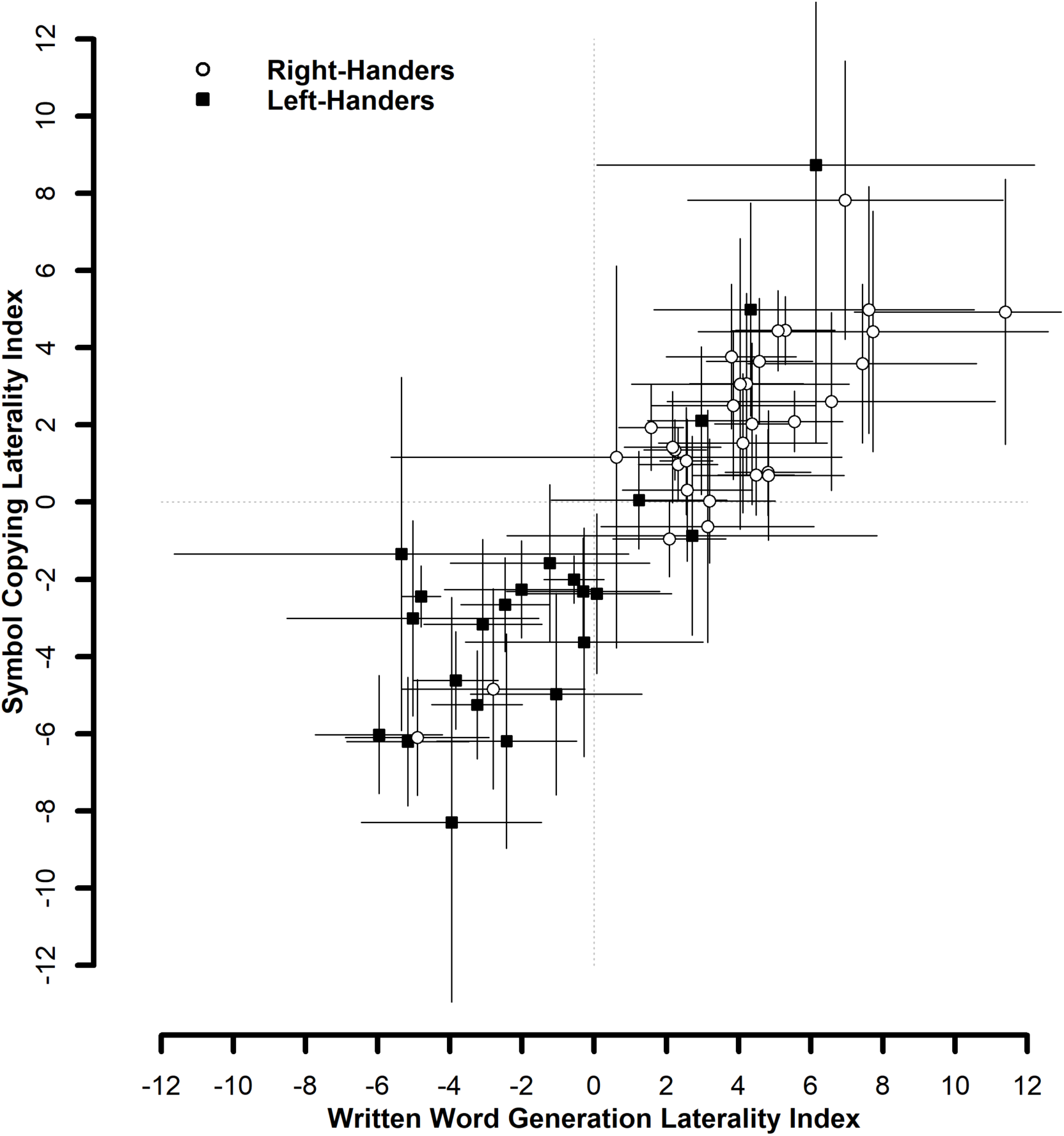
Scatter plot of laterality indices (LIs) for the written word generation and symbol copying conditions. All individual data points are presented with their 95% confidence intervals. Handedness groups according to the writing hand (right-handers: white circles; left-handers black squares).

To address the secondary aim, the relationships between LI_difference_ and the handedness measures are presented in Table 3. The relationships range from small to medium, but relatively consistent, which is unsurprising given the medium to large correlation between measures. The correlations for left- and right-handers (based on writing hand) are presented separately (see Table 3). Except for the relationship between EHI and QHPT in right-handers, the within-group relationships are heavily reduced in size, likely reflecting that the left- and right-handers provide variance along the continuum of hand preference, not varying qualitatively between groups.

**Table 3.** Spearman correlation coefficients for relationships between LI_difference_ and handedness measures

| Group |  | LI | EHI | Pegboard | QHPT |
| --- | --- | --- | --- | --- | --- |
| Overall | EHI | 0.4* |  |  |  |
|  | Pegboard | -0.26 | -0.69*** |  |  |
|  | QHPT | 0.38* | 0.57*** | -0.41* |  |
|  | Writing | -0.35* | -0.83*** | 0.77*** | -0.41** |
| Left-Handers | EHI | 0.27 |  |  |  |
|  | Pegboard | 0.15 | 0.05 |  |  |
|  | QHPT | -0.05 | 0.24 | -0.45 |  |
| Right-Handers | EHI | 0.17 |  |  |  |
|  | Pegboard | -0.11 | -0.27 |  |  |
|  | QHPT | 0.37 | 0.55* | -0.14 |  |

\* $p < .05$ , \*\* $p < .01$ , \*\*\* $p < .001$

## Discussion

The present study used functional transcranial Doppler ultrasound (fTCD) to compare left- and right-handers in a written word generation task with a symbol copying task: similar with respect to visuomotor demands but excluding language. Our main objective was to investigate whether the broader right-hemispheric network previously observed in left-handers compared to right-handers during written as opposed to oral language production (Kondyli et al., 2017) subserves linguistic or motor demands of writing.

Our hypothesis, namely that left-handers will exhibit weaker typical (left) linguistic-specific lateralization for written language generation compared to right-handers, was supported. We tested this hypothesis in two different ways: (i) By examining whether the LI_difference_ is different to zero for the two handedness groups. We found that the LI_difference_ was different to zero for right-handers, but not for left-handers; (ii) By examining whether left- and right-handers differ on the difference between written word generation and symbol copying. We indeed found a smaller LI_difference_ in left-handers compared to right-handers, as expected. Overall, right-handers presented with a more left-hemispheric activation pattern when isolating the linguistic aspect of writing, similarly to oral language tasks.

To explain the difference between right- and left-handed participants we explored whether (i) group variability, (ii) individual inter-trial LI_difference_ variability, (ii) the number of left-lateralized individuals differed between groups, and (iv) whether lateralization is weaker in left-lateralized left-handers compared to left-lateralized right-handers. No evidence of differences was found for any of these analyses. Not finding weaker lateralisation in left-lateralized left-handers contrasts with a recent finding that left-handers who are left-hemisphere dominant for language (about 70% of left-handers) are less lateralized compared to their right-handed counterparts (Johnstone et al., 2021). This difference may be attributed to the methods used, but also due to power (*n* = 40 for this analysis compared to *n* = 74 in Johnstone et al.). Another possibility is that the areas subserving language are broader in left-compared to right-handers, as suggested by Kondyli et al. (2017).

Given that the right-channel activation was stronger in the symbol copying condition only for right-handers, an alternate explanation is a contribution of a right-hemispheric attentional network for the more-novel stimuli (i.e., the symbols), similar to what has been observed in visuospatial tasks (Rosch et al., 2012; Whitehouse & Bishop, 2009). This contribution was not apparent in left-handers; however, the written word generation task already demanded the activation of right-hemispheric areas to support the motor demands of writing, therefore the activation related to the symbol copying task, which had the same motor demands in addition to the attentional demands, potentially reached a ceiling (see Figure 1). Therefore, fTCD may simply have been insensitive to this effect in left-handers. Future work could disentangle this by comparing copying of simple versus complex novel stimuli or over-learned nonverbal stimuli, such as shapes. Future work could also compare writing with letter copying. This will be especially important to understand patterns in left-handers who did not show clear differentiation between written word generation and symbol copying.

Another explanation is a graded increase during handwriting in functional activation in areas of the right hemisphere, namely the right anterior supramarginal gyrus, with the degree of left-handedness (Siebner et al., 2002). These areas could respond to motor elements of writing, such as motor preparation before handwriting, with left-handers possibly having more difficulty with task initiation and hence showing greater effort related to movement preparation (Siebner et al., 2000). Of note, this suggestion is based on functional imaging studies on right-handers that pointed towards a role in movement preparation and selection for the left inferior parietal lobule (Deiber et al., 1996; Krams et al., 1998; Schluter et al., 2001).

Previous evidence from studies comparing handwriting to similar motor tasks, such as clock drawing or the drawing of simple geometric shapes or objects, has shown that frontoparietal networks are activated. These networks include the superior parietal cortex, the supplementary motor area, the dorsal premotor and ventral premotor cortices, and the cerebellum (Ferber, Mraz, Baker, & Graham, 2007; Gowen & Miall, 2006; Ino, Asada, Ito, Kimura, & Fukuyama, 2003; Makuuchi, Kaminaga, & Sugishita, 2003; Miall, Gowen, & Tchalenko, 2009). However, Planton et al. (2017) found that the distinctive feature between a non-linguistic motor task (drawing) and writing in a sample of right-handers is the left-lateralization pattern of the graphemic/motor frontal area. This left-lateralization pattern was replicated here in right-handers, but not for left-handers.

A secondary aim of this study was to examine the relationship of a comprehensive set of handedness assessment criteria to cerebral lateralization estimates. As anticipated, hand preference measures had higher correlations with the LI_difference_ compared to the hand skill measure. In fact, there was no evidence of a correlation between the pegboard task and LI_difference_, as opposed to all other handedness measures. It could be argued that it is hand preference and not hand skill measures that are informative for cerebral laterality for writing. These results add to previous findings that point to the direction of treating hand skill and hand preference as two rather distinct concepts. For example, hand skill and hand preference have been suggested to be independently lateralized (Triggs et al., 2000). Another possibility is that the pegboard task as a measure of hand skill was not sensitive enough to capture the handedness effect. Indeed, different measures of hand skill have been found to have low correlations with each other (0.08-0.3), suggesting that they correspond to different dimensions of laterality and cannot be used interchangeably (Buenaventura Castillo, Lynch, & Paracchini, 2019).

On a methodological level, these findings showcase why it is important to measure and report handedness using more than one measure, as recently suggested by Papadatou-Pastou et al. (2020). The fact that handedness studies may use different measures of handedness and different criteria to group participants, has been highlighted repeatedly in recent meta-analyses as introducing noise to the literature, creating an obstacle to cross-study comparisons (Markou, Ahtam, & Papadatou-Pastou, 2017; Ntolka & Papadatou-Pastou, 2018; Papadatou-Pastou, Martin, Munafo, & Jones, 2008; Papadatou-Pastou et al., 2019; Papadatou-Pastou & Tomprou, 2015). In the fTCD literature, recently Kondyli et al. (2017) reported findings using the same four measures that were reported here and we urge researchers to adopt this good practice.

We shall refrain from making recommendations as to which measure is preferred, because each one has its own merits. Writing hand is the easiest, most intuitive, and popular method to access hand preference (Papadatou-Pastou et al., 2020). EHI is the most popular hand preference inventory in the literature (Papadatou-Pastou et al., 2020) and thus lends itself to cross-study comparisons. Moreover, it is easily administered in group settings or even online. The pegboard task is a measure of hand skill, therefore representing an important dimension of handedness that could be –as mentioned above-independently lateralized from preference. The QHPT measures preference behaviorally using an activity (card-reaching in different locations) that is not typically practiced in everyday life and is thus not expected to be subject to cultural pressures. Moreover, using the QHPT, preference can be more readily quantified than when using an inventory that lists different, unrelated activities. The pegboard and QHPT are administered physically on a one-to-one basis, making data collection more demanding. Considering these different merits and properties of the four handedness measures, we urge researchers to use and report as many of these methods as practically possible so that a better understanding of the multifaceted phenomenon of handedness can be achieved.

A potential limitation of the study is that the fTCD detects the blood flow in the MCAs, which feed mainly frontal and temporal brain areas (van der Zwan & Hillen, 1991; van der Zwan et al., 1993). While temporal areas have been associated with the linguistic component of writing, motor areas are found in the frontal lobe. Therefore, it could be the case that activation of parietal areas previously found to be important for writing, such as the left intraparietal sulcus and the left superior parietal area (Planton et al., 2017), might be missed by fTCD. However, more recent findings show that the MCA territory is more extensive than previously described, occupying approximately 54% of the supratentorial parenchymal brain volume and including the intraparietal sulcus (Kim et al., 2019). Still, neuroimaging techniques such as fMRI and PET could be employed in future work to complement the present findings with measurements of better spatial resolution. Another limitation of the present study could be the fact that the length of the writing period (35 seconds) did not allow for the exploration of early vs. late phase effects, which have been previously shown in written phonemic fluency tasks using fMRI (Golestanirad et al., 2015). These effects have not been investigated to date using fTCD or in relation to handedness, so future work could explore these areas. In addition to the above limitations, and while the experimental (i.e., the written word generation) and the control (i.e., the symbol copying) tasks were chosen as they share visuomotor but not linguistic demands, it could be said they further differ in the pace in which they are executed, with the experimental task possibly involving more pauses. Future studies could also compare left-handers who are left-hemisphere dominant to left-handers who are right-hemisphere dominant for oral language.

## Conclusions

By comparing a visuomotor task that includes a linguistic component (written word generation) with a task that has similar visuomotor demands without a linguistic component (symbol copying) we were able to show that the linguistic aspect of writing results in left-hemispheric activation similarly to the case of oral language production tasks in right-handers. It is potentially right-hemispheric language areas that are more engaged in left-handers and not merely motor areas, although attentional demands of symbol copying and/or visuomotor control more generally could also be at play.

## Data availability statement

All PsychoPy scripts, datasets, and analysis scripts have been uploaded to the Open Science Framework repository (https://osf.io/t79jh/)

## Author contributions

MPP and NB conceived and designed the study, MPP wrote draft manuscript, MPP and NB analyzed and interpreted findings, NB created the tables and figures, PS, AT, DS, IK, and SS collected data and assisted in data handling and analysis, MPP supervised data collection. All authors reviewed and approved the final version of the manuscript.

## Ethical statement

The study followed the declaration of Helsinki and has received ethical approval from the local ethics committee. All participants gave written informed consent.

## Abbreviations list

fTCD: functional transcranial Doppler ultrasound
fMRI: functional magnetic resonance imaging
PET: positron emission tomography
MCA: middle cerebral artery
QHPT: quantification of hand preference test
LI: laterality index
EHI: Edinburgh handedness inventory
POI: period of interest

